# Deploying a novel tuberculosis molecular bacterial load assay to assess the elimination rate of *Mycobacterium tuberculosis* in patients with multidrug-resistant tuberculosis in Tanzania

**DOI:** 10.1101/2020.09.02.280511

**Authors:** Peter M. Mbelele, Emmanuel A. Mpolya, Elingarami Sauli, Bariki Mtafya, Nyanda E. Ntinginya, Kennedy K. Addo, Katharina Kreppel, Sayoki Mfinanga, Patrick P.J. Phillips, Stephen H. Gillespie, Scott K. Heysell, Wilber Sabiiti, Stellah G. Mpagama

**Author notes:** **Corresponding author,** Dr. Peter Mbelele, Kibong’oto Infectious Diseases Hospital (KIDH), P.O BOX 12, Siha, Kilimanjaro, Tanzania.

## Abstract

**Background:** Rifampin or multidrug-resistant-tuberculosis (RR/MDR-TB) treatment has transitioned to injectable-free regimens. We tested whether *M. tuberculosis* (*Mtb*) elimination rates measured by molecular bacterial load assay (TB-MBLA) in sputa correlate with composition of the RR/MDR-TB antibiotic regimen.

**Methods:** Serial sputa were collected from patients with RR/MDR- and drug-sensitive TB at day 0, 3, 7, 14, and then monthly for 4 months of anti-TB treatment. TB-MBLA was used to quantify viable *Mtb* 16S rRNA in sputum for estimation of colony-forming-unit per mL (eCFU/mL). *Mtb* elimination rates were compared among regimens using nonlinear-mixed-effects modeling of repeated measures.

**Results:** Among 37 patients with a total of 296 serial sputa; 7 patients received rifampin/isoniazid/pyrazinamide/ethambutol (RHZE), 8 an all-oral bedaquiline-based regimen, 9 an injectable and bedaquiline-containing regimen, and 13 an injectable-containing but bedaquiline-free regimen. The overall mean daily *Mtb* elimination was −0.24 [95% Confidence-Interval (CI); −0.39 to −0.08)] log_10_ eCFU/mL, and it varied with treatment-regimen (p < 0.001). Compared to the adjusted *Mtb* elimination of −0.17 (95% CI; −0.23 to −0.12) for the injectable-containing but bedaquiline-free reference regimen, the elimination rates were −0.62 (95% CI; −1.05 to −0.20) log_10_ eCFU/mL for the injectable and bedaquiline-containing regimen (p = 0.019), −0.35 (95% CI; −0.65 to −0.13) log_10_ eCFU/mL for the all-oral bedaquiline-based regimen (p = 0.054), and −0.29 (95% CI; −0.78 to +0.22) log_10_ eCFU/mL for RHZE (p = 0.332)

**Conclusion:** TB-MBLA distinguished *Mtb* elimination rates in sputa from patients receiving different treatment regimens, suggesting a reliable monitoring tool for RR/MDR-TB, that does not require mycobacterial culture.

## Introduction

Measurement of pulmonary tuberculosis (PTB) treatment response in endemic settings largely depends on sputum smear microscopy ^[1]^. While the sputum smear microscopy detection threshold is at least 10^3^ *Mycobacterium tuberculosis* (*Mtb*) in colony-forming-units in 1 mL (CFU/mL) per sputum sample, many patients with PTB such as those with human immunodeficiency virus and the acquired immunodeficiency syndrome (HIV/AIDS) present with paucibacillary disease and may be unable to produce a good quality sputa for detection of acid-fast-bacilli (AFB) ^[2,3]^. Besides, sputum smear microscopy cannot differentiate drug susceptibility, thus it is not applicable for rifampicin and or multidrug resistant (RR/MDR)-TB diagnosis or treatment monitoring. Furthermore, microscopy cannot distinguish viable from non-viable *Mtb* which requires prolonged incubation in solid or liquid media ^[3]^. Patients with RR/MDR-TB are typically monitored for cultured growth in Lowenstein-Jensen (LJ) solid medium or the Mycobacterium Growth Indicator Tube liquid culture system. Culture is sensitive with a detection limit of 10 – 100 CFU/mL of sputum, yet it is also prone to contamination and can take up to 8 weeks to determine a definitive positive or negative result, thereby limiting the ability to take appropriate and timely clinical action ^[4]^.

The novel TB molecular bacterial load assay (TB-MBLA) was developed by Gillespie et al and used for monitoring clearance of *Mtb* from sputa, as a marker for TB treatment response ^[5]^. TB-MBLA is a real-time polymerase chain reaction (RT-qPCR) assay which detects and quantifies elimination of 16S rRNA from both viable replicating and dormant *Mtb* in patient’s sputa during treatment ^[6]^. Previously, TB-MBLA was assessed by the Pan-African Consortium for Evaluation of Anti-TB Antibiotics (PanACEA) group in patients treated for drug-sensitive (DS)-TB, and demonstrated considerable potential to replace both smear microscopy and culture for monitoring TB treatment response ^[6–8]^. TB-MBLA was found to be consistently read as positive for samples with as low as 10 CFU/mL of *M. tuberculosis* and the cycle threshold for this read-out has been optimized at a value of 30 ^[6]^.

Recently, TB-endemic countries, including Tanzania, have adopted new and repurposed TB medicines, such as bedaquiline, delamanid and linezolid, and constructed regimens with limited microbiological evidence of effectiveness in patients with RR/MDR-TB. Hence, we deployed TB-MBLA to describe elimination of *Mtb* in patients receiving RR/MDR-TB and DS-TB treatment. We tested the hypothesis that *Mtb* elimination rates from the sputa, as measured by TB-MBLA, not only correlated with time-to-culture conversion but were dependent upon the composition of the RR/MDR-TB antibiotic regimen.

## Materials and Methods

### Patients, ethics and design

From August 2018 to December 2019, longitudinal cohort study was conducted among patients with RR/MDR- and DS-TB confirmed using Xpert® MTB /Rif ^[9]^. The study was approved by the National Institute for Medical Research (NIMR) in Tanzania (NIMR/HQ/R.8a/Vol. IX/2662). Permission to conduct the study was granted by authorities of the Kibong’oto Infectious Diseases Hospital (KIDH). Inclusion criteria were patients aged at least 18 years who consented to provide quality early-morning sputum and clinical information. Critically ill patients, pregnant women and those who interrupted treatment were excluded. Each patient was followed for 16 weeks during which they provided sputum for testing at day 0 (baseline), 3, 7, 14, 28, 56, 84 and 112 of treatment. The treatment regimens included standard RHZE (rifampicin, isoniazid, pyrazinamide, ethambutol) for DS-TB; an all-oral bedaquiline based regimen (bedaquiline, linezolid, levofloxacin, pyrazinamide and ethionamide), an injectable and bedaquiline containing regimen (kanamycin, bedaquiline, levofloxacin, pyrazinamide and ethionamide), and injectable-containing but bedaquiline free regimen (kanamycin, levofloxacin, pyrazinamide, ethionamide and cycloserine) containing regimens for RR/MDR-TB.

### Study Setting

Patients were recruited at KIDH, national centre of excellence for clinical management of drug resistant (DR)-TB located in the Siha district of Kilimanjaro region in Tanzania ^[9]^. TB-MBLA testing was performed at the National Institute for Medical Research, Mbeya Medical Research Centre branch, given that laboratory’s prior experience with the assay.

### Sample size determination

The numbers of patients required to determine differences in bactericidal activity over time in 4 treatment regimens were calculated as previously reported by Guo et al ^[10]^. We assumed a Spearman correlation of 0.51, and a baseline *Mtb* burden of 5.5 log_10_ eCFU/mL, as well as daily *Mtb* decline and decay rate of 0.42 and 0.05 log_10_ eCFU/mL respectively ^[6,8]^. Hence, at least 9 patients were needed per regimen to reach a power of 90% with a two-sided type I error of 5%. Considering a RR/MDR-TB treatment success of 56% globally and 75% in Tanzania ^[11]^, at least 20% of patients were likely to lost be to follow up and hence a minimum of 45 patients were desirable to be sampled.

### TB-MBLA and Culture

*M. tuberculosis* quantification by TB-MBLA was performed as described by Gillespie et al ^[11]^. In summary, 1mL of homogenized sputum was treated using guanidine thiocyanate (GTC), and was frozen at −80°C to preserve the *M. tuberculosis* RNA. Total *M. tuberculosis* RNA was extracted using the RNA pro (FastRNA Pro BlueKit MP Biomedical) according to manufacturer’s instructions. The extract was treated with DNase I enzyme (TURBO DNA-Free Kit Ambion) to remove DNA. The *M. tuberculosis 16*S rRNA was quantified by reverse transcriptase quantitative PCR (RT-qPCR) and the cycle-threshold CT translated to bacterial load (estimated CFU per mL (eCFU/mL) using a standard curve on a Rotor gene Q 5plex platform (Qiagen). The cut-off for TB-MBLA positivity is a 30 CT value that corresponds to 1.0 log_10_ eCFU/mL, beyond which the test was considered negative ^[8,11]^. *Mtb* culture was performed on LJ slants from the remaining sputum collected at baseline, 14 days then monthly for 4 months per previous instructions ^[13]^.

### Statistical analysis

Data were recorded in a clinical case report form (CRF), entered and cleaned before statistical analysis. Patients who completed 8 treatment visits and had positive pre-treatment TB-MBLA results were analysed and visualised in R, version 4.0.2 (http://www.R-project.org). Continuous variables such as age, body-mass-index (BMI) in Kg/m^2^ and time to TB-MBLA negativity were described as median with their 25^th^ and 75^th^ interquartile range (IQR), and were compared using a Kruskal–Wallis test. Accordingly, proportions for HIV, gender, cavitary-disease and previous TB treatment were compared across different regimens using Chi-Square or Fischer’s exact test. The rate of *Mtb* elimination (log_10_eCFU/mL) was fitted on quadratic polynomial nonlinear-mixed-effects (NLME) for repeated measures as previous ^[14]^, using Baseline bacterial load, cavity, HIV, silicosis and gender as fixed effects. Individual patients were accounted for random effect. A model was reliably selected if had low Akaike-information-criterion but high intraclass-correlation-coefficient (Table 2). Effect size in mean *Mtb* load between two treatment regimens at month 4 were compared using one-way analysis-of-variance (ANOVA) and Tukey’s test for repeated measures ^[15]^. The median time to TB-MBLA and culture conversion to negative was estimated using the Kaplan-Meier method, and was compared across different regimens using a log-rank test ^[16]^. Cox Proportional-Hazards regression models were used to estimate the hazard ratios (HR) for *Mtb* elimination, and was adjusted for the effects of HIV, baseline bacillary load, cavitary disease, silicosis, gender, prior history of treatment for drug sensitive TB and clearance rate. The mean *Mtb* load at baseline was the cut-off that beyond 4.0 log_10_ eCFU/mL was considered as high bacterial load. Mean clearance was considered as high if it was above the overall mean clearance rate and low if it was below. Similarly, the overall mean rate of *Mtb* clearance per day was used as the cut-off for low and high rate of clearance. A p value < 0.05 was considered significance. A 95% confidence interval (CI) of the mean clearance rate and HR was included.

**Table 1.** Socio-demographic and clinical characteristics of patients per treatment regimen.

| Variable | All | RHZE<br>(n = 7) | Injectable ±<br>BDQ (n = 21) | All-oral BDQ<br>(n = 9) | p-value |
| --- | --- | --- | --- | --- | --- |
| Median age (IQR) | 37 (32 – 49) | 30 (29 – 33) | 42 (34-54) | 36 (33- 44) | 0.038 |
| Male (%) | 27 (73) | 4 (57) | 18 (86) | 5 (56) | 0.125 |
| Chest cavity, n (%) | 29 (78) | 7 (100) | 14 (67) | 8 (89) | 0.163 |
| Probable TB, n (%) | 34 (92) | 7 (100) | 18 (86) | 9 (100) | 0.568 |
| HIV positive, n (%) | 11 (20) | 0 (0) | 3 (14) | 8 (89) | 0.001 |
| TB/Silicosis, n (%) | 7 (19) | 1 (14) | 4 (19) | 2 (22) | 0.731 |
| Malnourished, n (%) | 22 (59) | 4 (57) | 11 (52) | 7 (78) | 0.432 |
| Retreatment, n (%) | 23 (62) | 5 (71) | 14 (67) | 4 (44) | 0.528 |
| Median BMI (IQR) | 18 (15 – 19) | 17 (15 – 20) | 18 (16 – 20) | 17 (15 – 18) | 0.301 |
| Median days spent<br>before care (IQR) | 84 (60 – 196) | 85 (68 – 93) | 84 (56 – 196) | 88 (68 – 365) | 0.778 |

BDQ, bedaquiline; BMI, body-mass-index; injectable± BDQ, kanamycin with or without BDQ and IQR, interquartile range.

**Table 2.** Fitting and selection of a reliable polynomial nonlinear mixed effects model for repeated measures.

| Polynomial models<br>(degree) | Intercepts<br>(log10 eCFU/mL) | Intraclass correlation<br>coefficient (ICC) | Standard<br>deviation (SD) | Akaike information<br>criterion (AIC) | Likelihood<br>ratio test | p value |
| --- | --- | --- | --- | --- | --- | --- |
| Non-poly (model 1) | 3.00 | 0.54 | 0.81 | 722.89 | 1 vs. 2 | < 0.001 |
| Squared (model 2) | 2.99 | 0.63 | 0.67 | 634.63 |  |  |
| Cubic (model 3) | 3.00 | 0.65 | 0.63 | 611.59 | 2 vs.3 | < 0.001 |
| Quadratic (model 4) | 3.20 | 0.67 | 0.61 | 592.7 | 3 vs. 4 | < 0.001 |
| Pentadratic (model 5) | 2.89 | 0.68 | 0.60 | 588.58 | 4 vs. 5 | 0.020 |

Model 4 had the lowest AIC and within variability (SD) but high ICC values, the key selection criteria for a reliable model, and hence it was used to model *M. tuberculosis* elimination rates

## Results

### Population

Of 59 patients enrolled, 37 patients with a total of 296 serial sputa were analysed. Reasons for exclusion and patient’s distribution are outlined in Figure 1. In total, 30 (81%) and 7 (19%) of 37 patients analysed had RR/MDR-TB and DS-TB respectively. Clinical and demographics are presented in Table 1. Twenty-seven (73%) out of 37 patients were male. Their median (IQR) age was 37 (32 – 49) years. Patients who received standard RHZE treatment were younger than those who received RR/MDR-TB treatment regimens (p = 0.038). Also, 11 (30%) patients were living with HIV infections with a CD4 T cell count of 208 (95% CI; 144 – 272) cells/µL. More patients with HIV received an all-oral than injectable-based treatment regimen (p = 0.001).

**Figure 1.**
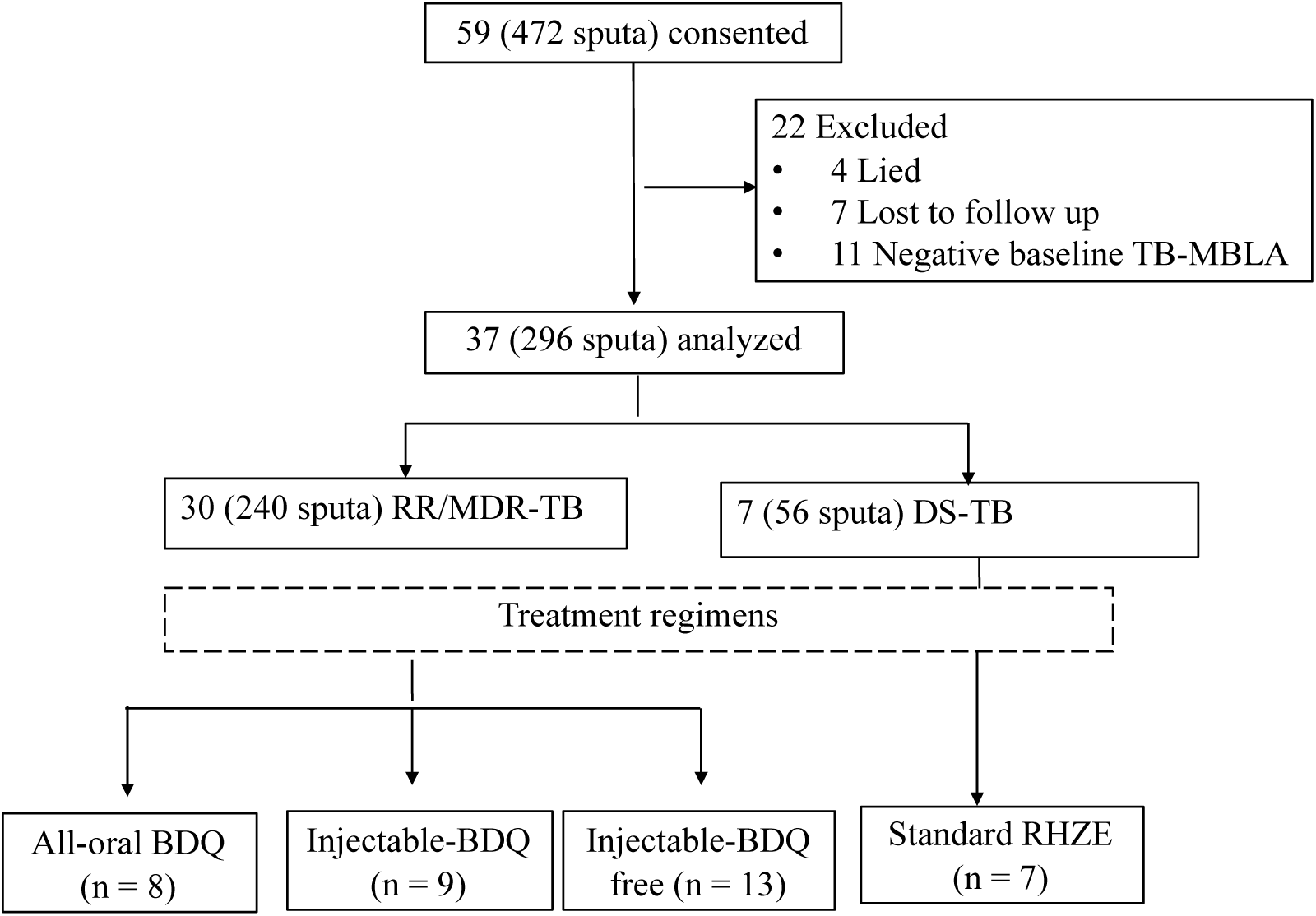
Recruitment and patient’s distributions in different treatment regimens. DS-TB, drug sensitive TB; RR/MDR-TB, rifampicin or multidrug resistance, TB-MBLA; tuberculosis molecular bacterial load assay, Standard RHZE comprised of rifampicin, isoniazid, PZA & ethambutol). Injectable-BDQ free regimen was comprised of kanamycin (KAN), levofloxacin (LFX), pyrazinamide (PZA), ethionamide (ETH) and Cycloserine (CS). Injectable-BDQ regimen was comprised of KAN, Bedaquiline (BDQ), LFX, PZA and ETH; and All-oral BDQ regimen contained BDQ, LFX, linezolid (LZD), PZA and ETH.

### Bactericidal activity over time

The *Mtb* load measured by TB-MBLA and culture in Figure 2 decreased significantly over time (R = −0.77, p < 0.001). The mean *Mtb* load in log_10_ eCFU/mL (95% CI) was reduced from 5.19 (4.40 – 5.78) at baseline to 3.10 (2.70 – 3.50) at day 14, then to 2.52 (2.13 – 2.90) at day 28, 1.88 (1.53 -2.22) at day 56 and <1.36 (1.03 – 1.70) at day 84 through 112 of treatment. The overall mean daily *Mtb* elimination was −0.24 (95% CI; −0.39 to −0.08) log_10_ eCFU/mL, and it varied with treatment-regimen (Table 3, p < 0.001). An injectable and bedaquiline containing regimen had the highest mean *Mtb* elimination rate followed by an all-oral bedauquiline based regimen compared to injectable-containing but bedaquiline free reference regimen (Table 3, p = 0.019). Kanamycin containing regimens in Figure 3 had rapid bactericidal activity at day 14, but was not translated into long term bactericidal effect (p < 0.001). An all-oral bedaquiline-based regimen had a sharp decline after day 28.

**Table 3.** Mean daily M. tuberculosis elimination rates (log_10_ eCFU/mL) and corresponding burden at day 0 (baseline) and 112 of treatment.

| Treatment regimens | Mean *M. tuberculosis* elimination rates |  |  |  | Mean (95% CI) *M. tuberculosis* load |  |
| --- | --- | --- | --- | --- | --- | --- |
|  | Unadjusted model for covariates |  | Adjusted model for covariates |  | Day 0 (baseline) <sup>†</sup> | Day 112 <sup>*</sup> |
|  | Rates (95% CI) | p-value | Rates (95% CI) | p-value |  |  |
| 1. Reference (injectable-BDQ free) | -0.18 (-0.27 to -0.08) |  | -0.17 (-0.23 to -0.12) |  | 4.73 (4.13 – 5.32) | 2.77 (2.51- 3.04) |
| 2. Injectable-bedaquiline | -0.48 (-1.25 to +0.28) | 0.239 | -0.62 (-1.05 to -0.20) | 0.019 | 4.63 (3.95 – 5.47) | 2.08 (1.81 - 2.36) |
| 3. All-oral bedaquiline | -0.26 (-0.48 to +1.00) | 0.507 | -0.35 (-0.65 to -0.13) | 0.054 | 5.36 (4.65 – 6.08) | 2.47 (2.20 - 2.74) |
| 4. Standard RHZE | -0.23 (-0.57 to +1.02) | 0.593 | -0.29 (-0.78 to +0.22) | 0.332 | 5.17 (4.36 – 5.99) | 2.51 (2.18 - 2.85) |

<sup>†</sup>Baseline mean *M. tuberculosis* load in all regimens were comparable (ANOVA, p = 0.453). An asterisk (\*) denotes p -values for mean difference in *M. tuberculosis* load for regimen pairwise comparison at day 112: regimen 1 & 2, p < 0.001; regimen 2 & 3, p = 0.031; regimen 1 & 3, p = 0.077; and regimen 2 & 4, p = 0.040. Reference regimen was the injectable-bedaquiline (BDQ) free regimen composed of kanamycin (KAN), levofloxacin (LFX), pyrazinamide (PZA), ethionamide (ETH) and Cycloserine (CS); Injectable-bedaquiline regimen was comprised of KAN, BDQ, LFX, PZA and ETH; All-oral bedaquiline regimen contained BDQ, LFX, linezolid (LZD), PZA and ETH; and the RHZE for rifampicin, isoniazid, PZA and ethambutol (E) Covariates adjusted included baseline bacterial load, cavity, gender, HIV and silicosis, *M. tuberculosis* elimination rates varied among regimens

**Figure 2.**
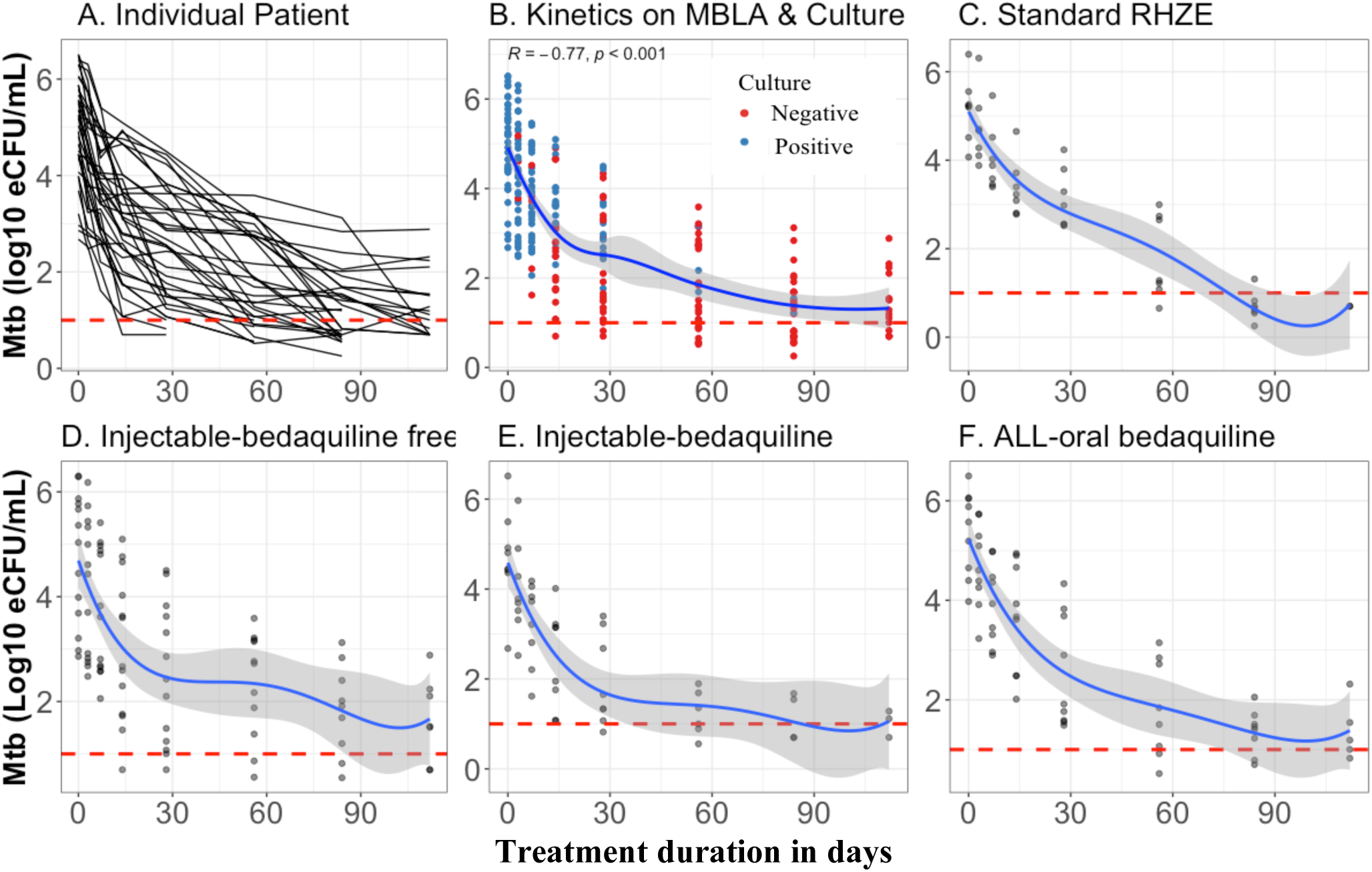
M. tuberculosis elimination during treatment. The plots A-F show *M. tuberculosis (Mtb)* kinetics between patients (A) as measured by TB-MBLA and culture (B) among patients treated with standard RHZE (C), injectable bedaquiline free regimen (D) containing kanamycin (KAN), levofloxacin (LFX), pyrazinamide (PZA), ethionamide (ETH) and Cycloserine (CS); Injectable-bedaquiline regimen (E) was comprised of KAN, Bedaquiline (BDQ), LFX, PZA and ETH; and an all-oral bedaquiline regimen (F) containing BDQ, LFX, linezolid (LZD), PZA and ET

**Figure 3.**
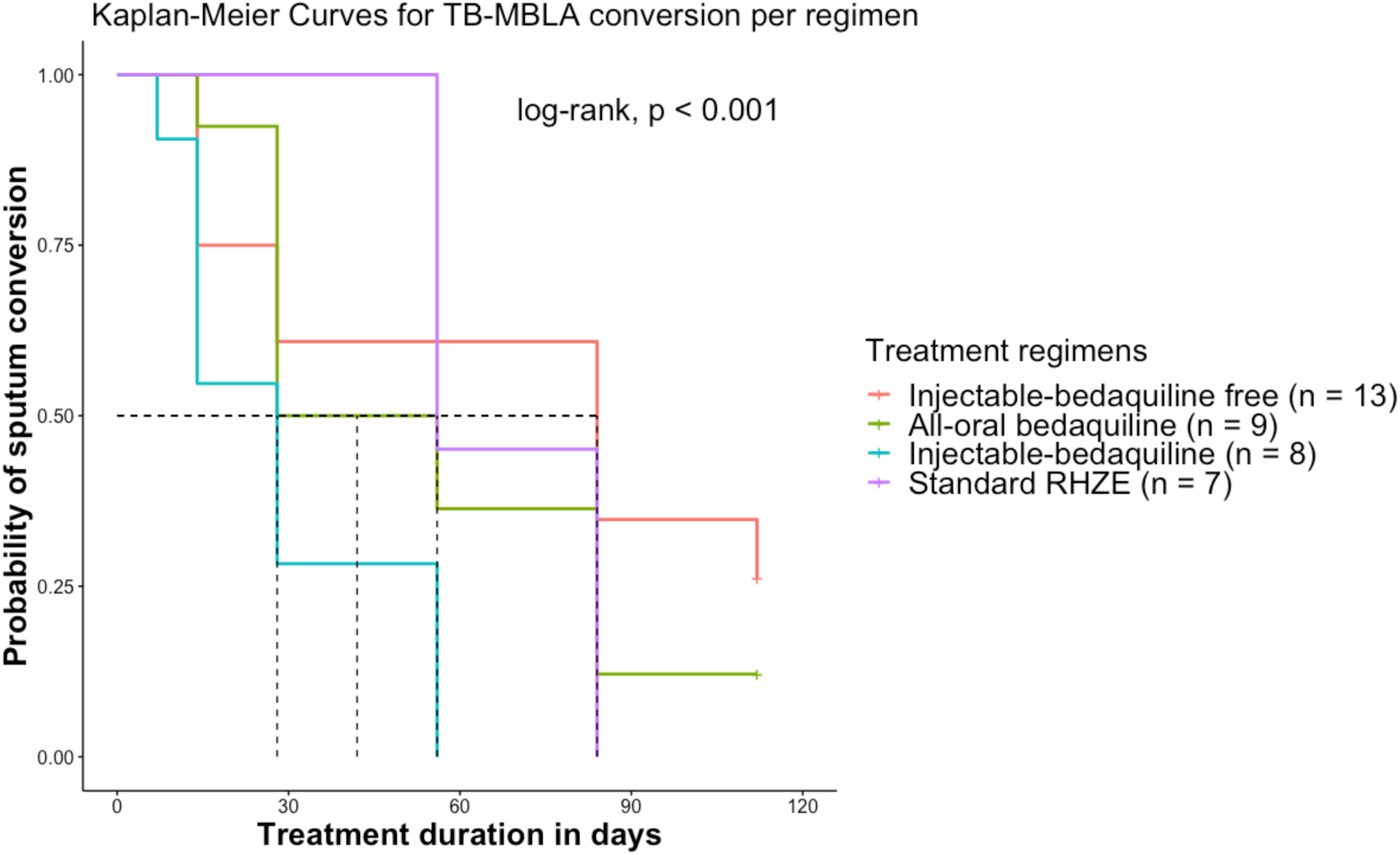
*M. tuberculosis* elimination per treatment regimen over time. Bedaquiline containing regimens had short median time to TB-MBLA conversion to negative compared to injectable-containing but bedaquiline free regimen containing kanamycin (KAN), levofloxacin (LFX), pyrazinamide (PZA), ethionamide (ETH) and Cycloserine (CS). Injectable-bedaquiline was comprised of KAN, bedaquiline (BDQ), LFX, PZA and ETH; an all-oral bedaquiline regimen was composed of BDQ, LFX, linezolid (LZD), PZA and ETH, and Standard RHZE composed of rifampicin, isoniazid, PZA and ethambutol.

### Median time to *M. tuberculosis* elimination

There was strong positive correlation in time-to sputum conversion between TB-MBLA and culture [r = 0.46 (95% CI; 0.36 – 0.55), p < 0.001]. The overall median time to sputum TB-MBLA conversion to negative was 56 (IQR; 28-84) days. The median time to TB-MBLA conversion to negative were 28, 42 and 84 days among patients on injectable and bedaquiline, an all-oral bedaquiline-based regimen, and injectable-containing but bedaquiline free regimens respectively. Percentage of patients who converted to sputum negative by TB-MBA and culture are shown in Figure 4. Approximately, 24% (9/37) of patients had negative TB-MBLA at day 14 compared to 51% (19/37) culture negative (p = 0.019), which was respectively increased to 43% (16/37) and 65% (24/37) at day 28 of treatment (p = 0.002). At day 56, 68% (25/37) had sputum converted to negative by TB-MBLA compared to 89% (33/37) by culture (p = 0.897). Despite that all patients on standard RHZE converted to negative at day 90 of treatment, 4 patients with RR/MDR-TB did not convert to negative. Three out of these 4 patients were on injectable-containing but bedaquiline-free, and remained positive by TB-MBLA at day 112

**Figure 4.**
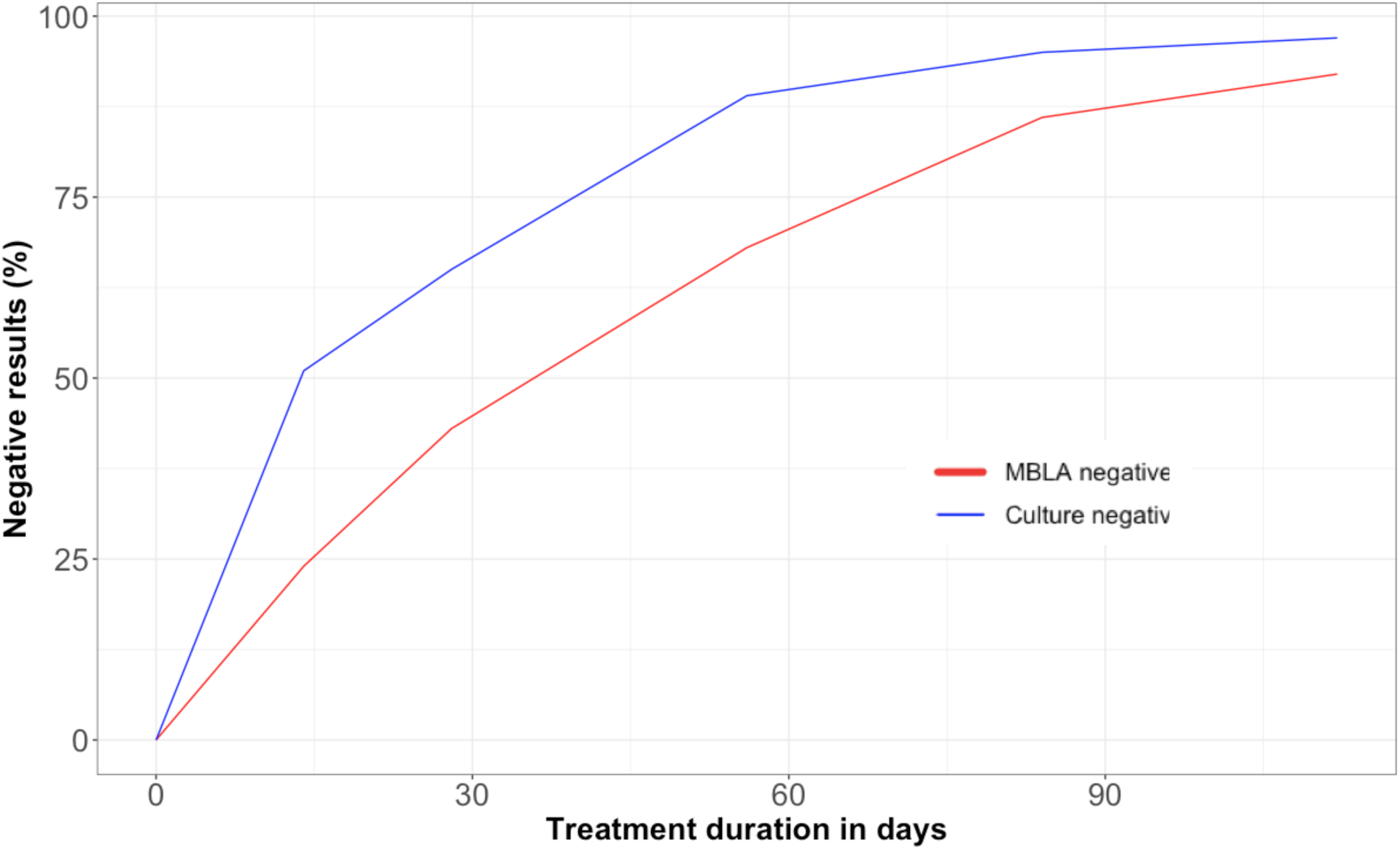
Percentage of patients who converted to negative by TB-MBLA and culture over time (N = 37). Changes in presentation of patients with negative results by tuberculosis molecular bacterial load assay (TB-MBLA, red line) and Lowenstein-Jensen culture medium (blue line). In the first 60 days, high proportion of patients had negative culture results compared to TB-MBLA.

### Hazard ratio (HR) of *M. tuberculosis* elimination

The overall mean Mtb load log_10_ eCFU/mL at baseline was 5.19 (95% CI; 4.40 – 5.78), and was similar in all patients treated with any of the 4 regimens (Table 3, p = 0.453). The mean Mtb load (log_10_ eCFU/mL) among female was 5.6 (95% CI; 5.0 – 6.2) log_10_ eCFU/mL compared to 4.7 (95% CI; 4.3 – 5.2) log_10_ eCFU/mL among male (p = 0.017) patients. Patients with chest cavity had mean Mtb load of 5.26 (95% CI; 4.45 – 5.87) compared to 4.40 (95% CI; 3.91 – 4.75) log_10_ eCFU/mL in those without cavity (p = 0.080). Adjusting for bacterial load, initial elimination rate, silicosis, chest cavity, HIV and gender, the hazard-ratios for *Mtb* elimination were 12.37 (95% CI, 2.87 – 53.30; p = 0.001) and 14.31 (95% CI, 3.49 – 58.65; p < 0.001) for patients who received an all-oral bedaquiline and injectable and bedaquiline-containing regimens respectively (Table 4). Bacterial load at baseline strongly correlated positively with median time to sputum conversion to negative by both TB-MBLA and culture [r = 0.48 (95%CI; 0.18 – 0.69), p = 0.003). High Mtb load and TB/silicosis were independently predictor of slow Mtb elimination compared to low Mtb load and TB without silicosis (Table 4, p ≤ 0.033

**Table 4.**
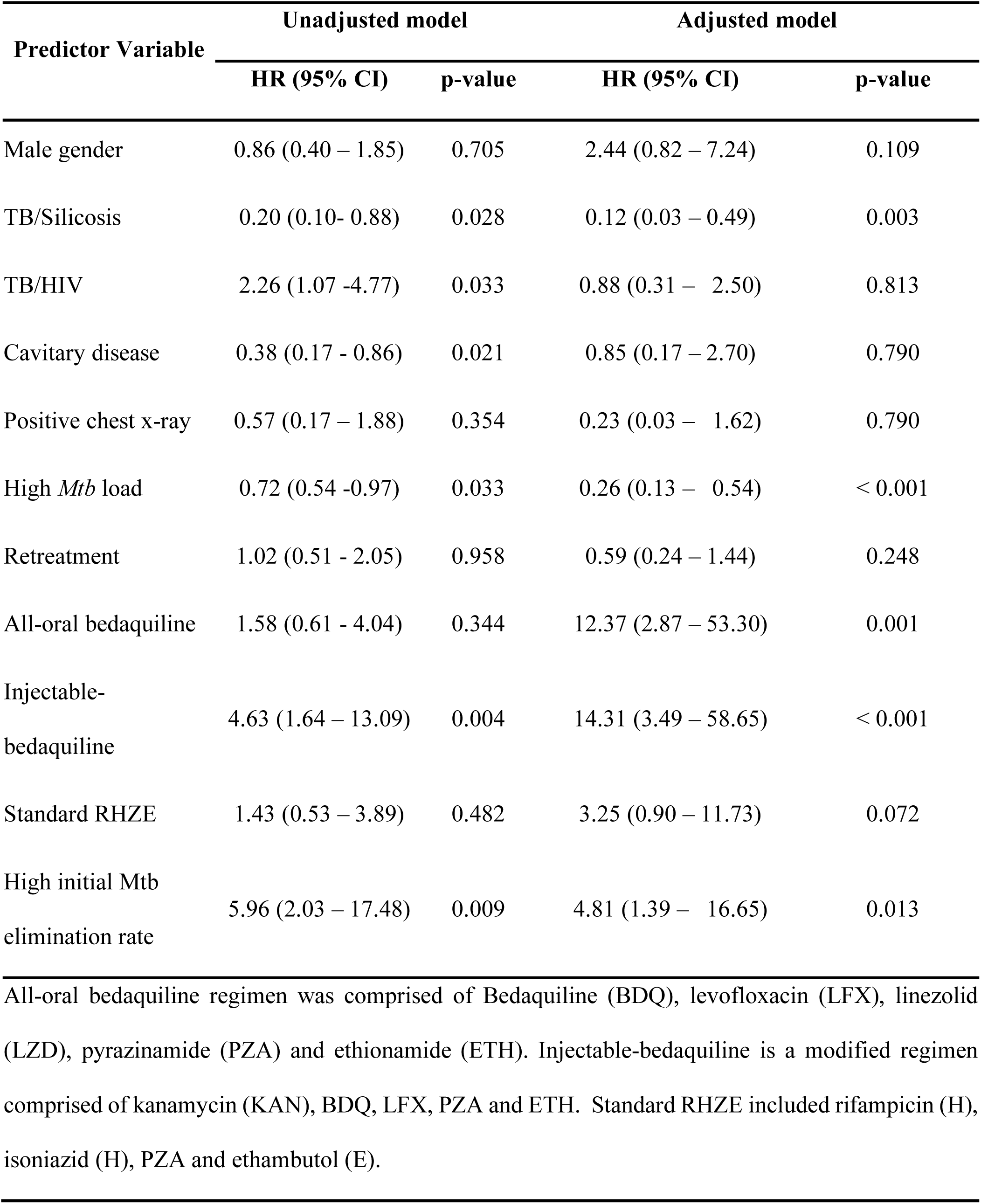
Hazard ratio (HR) of *M. tuberculosis* (*Mtb*) elimination in Cox Proportion-Hazard model.

## Discussion

This study shows for the first time to our knowledge that TB-MBLA is promising for monitoring treatment response among patients treated with DS- and -RR/MDR-TB regimens, as well as those with concomitant TB/silicosis. As measured by TB-MBLA, *M. tuberculosis* decreased significantly over time on treatment, and this kinetic correlated with what was observed using LJ culture medium. For decades, culture has been used as a routine microbiological tool for monitoring drug-resistant TB treatment response ^[17,18]^, but in many TB endemic settings, culture is unavailable or limited to specialized centres. Importantly, culture results can take up to 8 weeks from the time of sputum collection, which when making treatment decisions based on a result from a two-months old specimen, is akin to driving a car while only looking in the rear-view mirror. Given the continued decentralization of RR/MDR-TB services, monitoring treatment response in laboratories capable of performing qPCR, such as with Xpert MTB/RIF, will allow laboratory assays to impact treatment decisions closer to the point-of-care. Therefore this study in RR/MDR-TB compliments the growing evidence base for the application of TB-MBLA in routine clinical management ^[6,8,19]^.

Interestingly, our findings suggest that bactericidal activity at day 14 may not be a suitable predictor of the long-term efficacy of a regimen, particularly when that regimen is bedaquiline containing. In this cohort at day 14, more than 75% of people had a positive TB-MBLA and more than half had a positive culture result. Whereas between 14-56 days we observed substantial *M. tuberculosis* elimination in those treated with a bedaquiline containing regimens, suggesting that evaluation of bactericidal activity be performed later, such as at day 56, for modern RR/MDR-TB regimens. These findings may contradict those from a phase 2b trial where the bactericidal activity of a bedaquiline containing regimen as was measured by culture media at day 56 proved an unreliable indicator of a regimen’s ability to predict long term treatment outcomes or shorten treatment duration, and rather raise the question of whether TB-MBLA may in fact be a superior predictor to culture.^[20]^

Another important finding from this study of TB-MBLA is that *M. tuberculosis* elimination kinetics were regimen-dependent. Overall, more rapid elimination occurred during the first 28 days for all regimens, yet that earlier rapid elimination was more prominent at day 14 for patients who received kanamycin regardless of receipt of bedaquiline, followed by those who received an all-oral bedaquiline containing regimen, which did not achieve these rates of elimination until 1 month or more of treatment. This observation concurs with previous reports that the bactericidal activity of bedaquiline in MDR-TB is delayed at the beginning, but accelerates later in therapy ^[21]^. Despite the superior activity of kanamycin containing regimens at day 14, this more rapid early elimination of *M. tuberculosis* was not sustained as a long term-bactericidal effect, such that 3 patients on injectable containing but bedaquiline free regimen remained positive after 4 months of treatment. These findings as measured by TB-MBLA fit with the pharmacodynamical understanding that kanamycin and other aminoglycoside/polypeptides if active against mycobacteria, primarily exert their effect against those extracellular organisms that are rapidly dividing and may be more abundant early in the treatment course ^[22,23]^.

The shorter overall time to sputum conversion to negative, as measured by TB-MBLA and conventional culture, for all patients who received bedaquiline regardless of kanamycin further supports arguments that bedaquiline should be a cornerstone of regimens designed to shorten MDR-TB treatment duration ^[24]^. The conventional injectable-containing but bedaquiline free regimen has been in practice for decades, even though more than 40% of patients treated with this regimen had unfavourable outcomes in TB endemic settings ^[11]^. Aminoglycosides such as kanamycin is no longer part of the current MDR-TB treatment regimens not because of its lack of bactericidal activity, as our data would suggest the contrary in the early treatment period, but rather because of the significant toxicity and patient intolerances that led to treatment interruption ^[25,26]^. While we do not advocate this approach, from microbiological perspective alone, as demonstrated in this study and others such as Mpagama *et al*.^[27]^, kanamycin could be included for first month only for instance and then dropped before toxicities accumulate. In a more patient-centered approach however, our findings demonstrate how potentially important it will be to find tolerable substitutes for kanamycin that can match the early bactericidal effect.

The main strengths in this study is that we have utilized TB-MBLA to model elimination rates among patients with RR/MDR-TB and those with TB/silicosis. We have shown that patients with TB/silicosis had slower *M. tuberculosis* elimination rates by TB-MBLA compared to those with TB and without silicosis. This slow rate of elimination could partially be attributed to the underlying pulmonary pathophysiology which can include progressive massive fibrosis ^[28,29]^, and anatomically, a blunted local host immune response to *M. tuberculosis* infection ^[28]^. We observed a similarly slower rate of *M. tuberculosis* elimination among patients with RR/MDR-TB who had high initial bacterial load, which supplements previous studies of TB-MBLA kinetics from patients with drug sensitive TB ^[6,8,19]^. Limitations of the study include the endpoints, which were limited to 4 months such that predicting long-term treatment success was beyond the scope of this study. Nevertheless, modelling *M. tuberculosis* elimination for 4 months as we accomplished here has been used as marker for treatment failure and relapse in several observational studies ^[18,30]^, and exceeds the duration of monitoring used in other trials of R/MDR-TB regimens that have employed conventional culture based techniques. ^[20]^ Additionally, this study had no control over the treatment regimens prescribed. However, given the feasibility of TB-MBLA and the comparability of this study’s findings to those prior with TB-MBLA in drug-susceptible TB ^[8]^, we plan to apply TB-MBLA systematically within an ongoing operational research protocol for injectable-free RR/MDR-TB treatment in Tanzania, that employs standardized regimens over varying treatment durations. Lastly, the number of patients per treatment regimen were small such that findings should be cautiously interpreted with inference to other populations with RR/MDR-TB. However, considering the low MDR-TB burden in countries like Tanzania as well as the repeated measurements per patient, findings in this study are critical to inform how TB-MBLA may be applied as a culture-independent method for RR/MDR-TB care locally.

In conclusion, patients who received bedaquiline-containing regimens exhibited higher *M. tuberculosis* elimination-rates and had shorter time-to sputum TB-MBLA and culture conversion to negative. While both kanamycin containing regimens had superior bactericidal activity during two weeks of RR/MDR-TB treatment, the addition of bedaquiline allowed for improved elimination after 1 month of therapy. Together, these findings provide insight into formulating optimal all-oral bedaquiline containing regimens with the best potential to shorten MDR-TB treatment duration ^[20,26,31]^. Given the ease of use of TB-MBLA and the fact that it does not require laboratory procedures associated with culture or the prolonged time to receive a culture-based result, we envision that TB-MBLA can be used to make regimen adjustments, and enhance infection control practices for patients with RR/MDR-TB and health workers in hospital and community settings

## Acknowledgements

This study received financial support from the EDCTP2 programme supported by the European Union project (grant number: TMA2016SF-1463-REMODELTZ) and DELTAS Africa Initiative (Afrique One-ASPIRE /DEL-15-008). The Afrique One-ASPIRE is funded by a consortium of donors including the African Academy of Sciences, Alliance for Accelerating Excellence in Science in Africa, the New Partnership for Africa’s Development Planning and Coordinating Agency, the Wellcome Trust (107753/A/15/Z), and the UK Government. All funding bodies have had no role in the conceptualization, methodology, data interpretation and writing of manuscript.

Furthermore, authors acknowledge Ms Batuli Mono, Taji Mnzava, Joseph Kachala and Dr Bibie Said of KIDH for their assistant with recruitment and data collection from study participants. We also thank Mr. Elisha S. Juma and Ms Sarapia P. Malya of KIDH, and Emmanuel Sichone and Joseph John of NIMR Mbeya for assisting with laboratory work. In addition, we also acknowledge the KIDH administration for granting permission to conduct this study.

## Transparency declarations

All authors have no conflict of interest to declare. PMM, EAM, ES, WS and SGM conceived the study, designed the work and interpreted clinical and TB-MBLA results. PMM and BM acquired data. PMM, KK, EAM, PPJP, WS, and SGM analyzed the data. PMM drafted the manuscript and responded to all co-authors’ inputs. SHG, NEN, and SKH reviewed the manuscript. All authors wrote, approved and agreed to be accountable for all scientific aspects in the final version of this manuscript.

